# Optogenetic dissection of mitotic spindle positioning in vivo

**DOI:** 10.1101/319772

**Authors:** Lars-Eric Fielmich, Ruben Schmidt, Daniel J. Dickinson, Bob Goldstein, Anna A. Akhmanova, Sander J.L. van den Heuvel

## Abstract

The position of the mitotic spindle determines the plane of cell cleavage, and thereby the location, size, and content of daughter cells. Spindle positioning is driven by dynein-mediated pulling forces exerted on astral microtubules. This process requires an evolutionarily conserved complex of Gα-GDP, GPR-1/2^Pins/LGN^, and LIN-5^Mud/NuMA^ proteins. It remains unknown whether this complex merely forms a membrane anchor for dynein, or whether the individual components have additional functions, for instance through Gα-GTP or dynein activation. To functionally dissect this system, we developed a genetic strategy for light-controlled localization of endogenous proteins in *C. elegans* embryos. Controlled germline expression and membrane recruitment of the Gα regulators RIC-8^Ric-8A^ and RGS-7^Loco/RGS3^, and replacement of Gα with a light-inducible membrane anchor demonstrated that Gα-GTP signaling is dispensable for pulling force generation. In the absence of Gα, cortical recruitment of GPR-1/2 or LIN-5, but not dynein itself, induced high pulling forces. Local recruitment of LIN-5 overruled normal cell-cycle and polarity regulation, and provided experimental control over the spindle and cell cleavage plane. Our results define Gα∙GDP–GPR-1/2^Pins/LGN^ as a regulatable membrane anchor, and LIN-5^Mud/NuMA^ as a potent activator of dynein-dependent spindle positioning forces. This study also highlights the possibilities for optogenetic control of endogenous proteins within an animal system.

## INTRODUCTION

Animal cells control the position of the spindle to determine the plane of cell cleavage. Regulated spindle positioning is therefore critical for asymmetric cell division and tissue formation (Pietro et al., 2016). Early work in *C. elegans* demonstrated that cortical pulling forces position the spindle through a protein complex that consists of a heterotrimeric G protein alpha subunit, GOA-1^Gαo^ or GPA-16^Gαi^ (together referred to as Gα) (Gotta and Ahringer, 2001; Grill et al., 2001; Miller and Rand, 2000), the TPR-GoLoco domain protein GPR-1/2 (Colombo et al., 2003; Gotta et al., 2003; Srinivasan et al., 2003), and the coiled-coil protein LIN-5 (Lorson et al., 2000). This complex, and the closely related *Drosophila* Gα_i/o_-Pins-Mud (Bellaïche et al., 2001; Bowman et al., 2006; Izumi et al., 2004; Schaefer et al., 2001) and mammalian Gα_i/o_-LGN-NuMA (Du and Macara, 2004; Du et al., 2001; Zheng et al., 2010; Zhu et al., 2011), recruits the microtubule motor dynein to the cell cortex (Nguyen-Ngoc et al., 2007) (Fig. 1a). While regulation at the level of individual components has been described, it remains unclear whether these proteins only form a physical anchor for dynein, or whether individual subunits contribute additional functions in spindle positioning.

**Figure 1.**
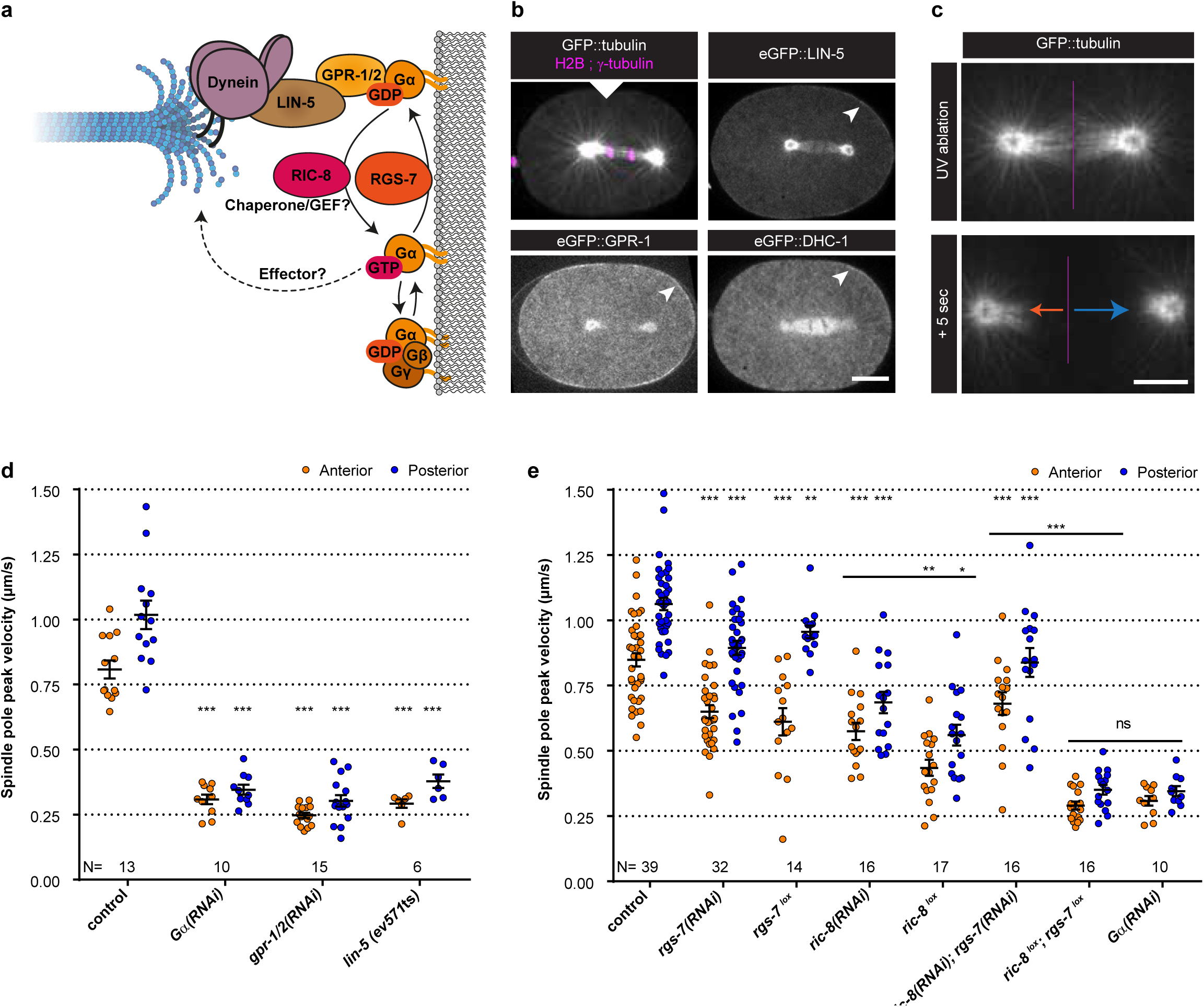
Gα regulaon by RIC-8 and RGS-7 is essenal for corcal pulling force generaon. **(a)** Cartoon model represenng mechanisms and funcons discussed in the text. Gα·GDP-GPR-1/2-LIN-5-dynein anchors dynamic microtubule plus-ends and generates corcal pulling forces on the mitoc spindle. Gα·GDP can assemble a Gαβγ or Gα-GPR-1/2-LIN-5 trimer. Gα·GDP/GTP nucleode state is regulated by the GAP RGS-7. For RIC-8, funcons as Gα GEF and chaperone are reported. Gα·GTP might promote spindle posioning through unknown downstream effectors. **(b)** Spinning disk confocal images of anaphase spindle posioning away from the cell center (white triangle) in the *C. elegans* zygote. The upper le panel shows the spindle with labeled tubulin and DNA. Other panels: endogenous GPR-1, LIN-5, and dynein (DHC-1) fused to eGFP are present in the cytoplasm, at the cortex (arrowheads), and spindle structures. Scale bar: 10 µm. **(c)** Spinning disk confocal images of the mitoc spindle (marked by GFP::tubulin). Upon UV-laser ablaon of the spindle midzone (violet line), spindle poles separate with velocies that represent the respecve net force acng on each pole (arrows). Scale bar: 5 µm. **(d)** Spindle pole peak velocies aer midzone ablaon. Control is *gfp::tubulin*. Other condions: inacvaon of Gα, GPR-1/2, and LIN-5. Error bars: s.e.m. Welch’s Student’s t-test; *** P < 0.001. **(e)** Spindle severing experiments in embryos where RIC-8 and RGS-7 were depleted by RNAi or induced ssue specific Cre-lox-mediated knockout of the endogenous gene (lox). Control is *gfp::tubulin*, see Supplementary Fig. 1 for knockout method and addional controls. Error bars: s.e.m. Welch’s Student’s t-test and Mann Whitney U test; * P < 0.05, ** P < 0.01, *** P < 0.001. See Supplementary Table 1 for detailed genotypes. Anterior is to the le in microscopy images.

As a potential additional function, force generation may require a dynein adaptor that activates dynein motility. Such an adaptor is necessary for the processive movement of mammalian cytoplasmic dynein during cargo transport along microtubules (Reck-Peterson et al., 2018). This process differs substantially form dynein-dependent cortical pulling, in which force is generated by shrinking microtubules (Laan et al., 2012). Without adaptor, surface-anchored yeast dynein in contact with depolymerizing microtubules generates pulling forces *in vitro* (Laan et al., 2012). However, yeast dynein moves processively on its own (Reck-Peterson et al., 2006). Hence, it remains unknown whether pulling force generation in animal cells depends just on anchoring of dynein, or whether this requires an additional dynein activator.

The role of Gα_i/o_ subunits in pulling force generation has also remained ambiguous (Fig. 1a). Membrane-attached Gα∙GDP associates with GoLoco motifs present in the homologous GPR-1/2, Pins, and LGN proteins (Kimple et al., 2002; Schaefer et al., 2001). This preference for the GDP-bound ‘inactive’ Gα state explains why RGS-7, a putative GTPase activating protein (GAP), promotes spindle positioning (Hess et al., 2004). However, the role of another conserved regulator of Gα signaling, RIC-8, remains poorly understood (Afshar et al., 2004; Miller and Rand, 2000; Tall et al., 2003). RIC-8 was shown to act as a guanine nucleotide exchange factor (GEF) *in vitro* (Afshar et al., 2004; Tall et al., 2003), while it may function in vivo as a Gα chaperone (Afshar et al., 2005; David et al., 2005; Gabay et al., 2011; Hampoelz et al., 2005; Wang et al., 2005), or as both a GEF and chaperone. In addition to RIC-8, G-protein coupled receptors and Gαo∙GTP signaling contribute to spindle positioning in *Drosophila* neuroblasts and sensory organ precursor cells (Katanaev et al., 2005; Schaefer et al., 2001; Yoshiura et al., 2012). Therefore, it has been proposed that the Gα GTP-binding and hydrolysis cycle forms a critical step in cortical pulling force generation (Afshar et al., 2004; Srinivasan et al., 2003; Yoshiura et al., 2012). However, it is difficult to distinguish whether Gαo∙GTP contributes to force generation, or more indirectly relays cell-cell signaling to the spindle.

Here, we describe an optogenetic strategy for the systematic examination of individual contributions of cortical pulling force components in vivo. We use the *C. elegans* one-cell embryo (P0), which undergoes reproducible spindle positioning and asymmetric cell division in the absence of cell-cell signaling (Supplementary Video 1) (Rose and Gönczy, 2014). We developed methods for knock-out of endogenous genes and expression of foreign sequences in the *C. elegans* germline. This allowed the light-controlled localization of endogenous proteins through ePDZ-LOV domain interactions in the early *C. elegans* embryo. Our results show that Gα∙GDP and GPR-1/2 can be replaced with a light-inducible membrane anchor, while LIN-5 is required as activator of dynein-dependent cortical pulling force generation.

## RESULTS

We set out to systematically investigate the individual roles of the proteins involved in cortical pulling force generation. Our previous studies and CRISPR/Cas9-assisted endogenous tagging demonstrated that cytoplasmic dynein and the Gα-GPR-1/2–LIN-5 complex overlap and function together in pulling force generation at the cell cortex of *C. elegans* early blastomeres (Fig. 1b) (Schmidt et al., 2017; van der Voet et al., 2009). As a read-out for pulling forces, we measured spindle pole peak velocities after UV-laser ablation of the spindle midzone (Grill et al., 2001) (Fig. 1c and Supplementary Video 2). Interfering with Gα, GPR-1/2, or LIN-5 function abolished significant force generation, as previously reported (Fig. 1d). RNA interference (RNAi) of *ric-8* or *rgs-7* by dsRNA injection resulted in partial loss of pulling forces (Fig. 1e). Double *ric-8(RNAi); rgs-7(RNAi)* did not further decrease pulling forces as might be expected when RIC-8 and RGS-7 both promote a critical GTPase cycle (Hess et al., 2004; Srinivasan et al., 2003). However, RNAi of *ric-8* and *rgs-*7 is known to cause incomplete gene inactivation, which could also explain the limited defects. We generated germline-inducible knock-out alleles to circumvent this caveat. We inserted *lox* sites in the endogenous *ric-8* and *rgs-7* loci by CRISPR/Cas9-assisted recombineering (Fig. S1a) (Dickinson et al., 2013), and expressed the CRE recombinase specifically in the germline (Fig. S1b). Compared to the control without CRE activity, knockout embryos showed reduced spindle pole peak velocities *(ric-8^lox^*: anterior −50% and posterior −48%; *rgs-7^lox^*: anterior −29% and posterior −11%), similar to or more defective than the corresponding RNAi embryos (Fig. 1e, Fig. S1c, d, e). Importantly, the double knock-out of *ric-8^lox^; rgs-7^lox^* showed much reduced spindle pole movements (anterior −68% and posterior −67 %), thereby resembling *Gα(RNAi)* (Fig. 1e). This indicates that RIC-8 and RGS-7 act independently, or partly redundant, as positive regulators of Gα.

To gain further insight into the individual functions of cortical pulling force regulators, we sought to obtain spatiotemporal control of protein localization. To this end, we explored implementing the ePDZ-LOV system, which makes use of exposure to blue light to control protein heterodimerization (Harterink et al., 2016; Strickland et al., 2012). As introduction of *epdz*, *lov*, and *cre* sequences induced strong germline silencing responses, we developed a computational approach to design protein-coding sequences that are resistant to silencing in the germline. Our design algorithm assembles a coding sequence for any desired polypeptide from a list of 12-nucleotide words found in native germline-expressed genes (Fig. S2a). We hypothesized that transgenes designed in this way would mimic native genes and thereby evade the germline silencing machinery. Indeed, using this approach, we could obtain robust expression of several foreign transgenes that were otherwise silenced (Fig. S2b, c). Although some of these transgenes were stably expressed for many generations, others showed gradual silencing when passaging strains in culture (Fig. S2d); therefore, as a further buffer against silencing, we combined our germline-optimized exons with poly-AT-cluster rich intron sequences, which were recently demonstrated to protect against germline silencing (Frøkjær-jensen et al., 2016; Zhang et al., 2018). This combined approach resulted in stable germline expression of transgenes and enabled implementation of the ePDZ-LOV system for use in the *C. elegans* early embryo.

To characterize the ePDZ-LOV system, we created a strain with a membrane-bound LOV2 domain, expressed as a pleckstrin-homology domain (PH)–eGFP protein fusion (PH::LOV), together with cytosolic ePDZ::mCherry (Fig. 2a). Illumination with a blue (491 nm) laser rapidly induced recruitment of ePDZ::mCherry to PH::LOV, and allowed both global and local cortical enrichment in one-cell embryos (Fig. 2b and Supplementary Videos 3–5). Note that GFP is also excited with blue light. Hence, experiments that involve GFP imaging imply global and continuous induction of the ePDZ-LOV interaction. To test the reversibility of the ePDZ-LOV interaction, we followed ePDZ::mCherry membrane localization after a global activation pulse, and found dissociation kinetics similar to those reported by others (Hallett et al., 2016) (t ½ = 42 s; Fig. 2c and Supplementary Video 6). Thus, we conclude that the ePDZ-LOV system is suitable for controlled protein localization in the early *C. elegans* embryo.

**Figure 2.**
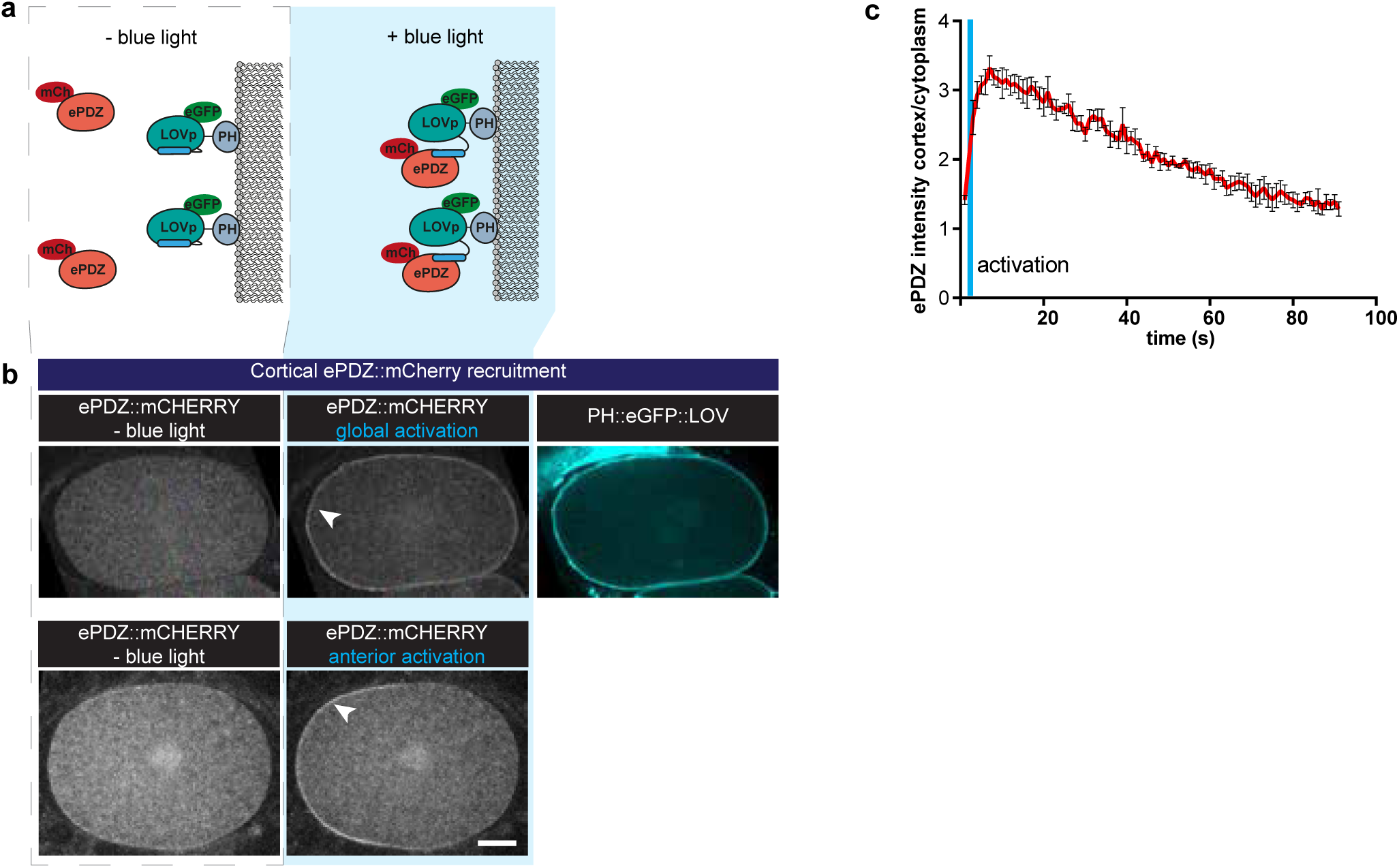
Opmized ePDZ-LOV enables light-inducible control of endogenous protein localizaon in the *C. elegans* one-cell embryo. **(a)** Cartoon model illustrang the proof of concept wherein cytosolic ePDZ::mCherry is corcally recruited to membrane PH::LOV upon acvation with blue light. **(b)** Spinning disk confocal images showing light-controlled localizaon of proteins in the *C. elegans* zygote (arrow-heads). Also see Supplementary Movies 3–5. Scale bar: 10 µm. **(c)** Quanficaon of corcal ePDZ::mCherry enrichment measured over me aer a 1 second pulse acvaon (blue vercal line). Error bars: s.e.m. t1/2 calculated with single component non-linear regression. Anterior is to the le in microscopy images.

Next, we examined whether membrane recruitment of RGS-7 and RIC-8 promotes pulling forces. Regulation of the Gα∙GDP/GTP cycle normally takes place at the cell membrane, while chaperoning of Gα folding and trafficking is expected to occur in the cytosol and at endomembranes (Gabay et al., 2011). We created strains expressing endogenous RIC-8 and RGS-7 as ePDZ::mCherry protein fusions. When combined with PH::LOV, this resulted in light-inducible membrane recruitment of RIC-8 and RGS-7 (Fig. 3a, d and Supplementary Video 7). Cortical enrichment of RGS-7 enhanced spindle pole movements (anterior +25% and posterior +20%) (Fig. 3b) and spindle oscillations (Fig. 3c). The RGS-7::ePDZ signal was too subtle to reliably control its local recruitment. As an alternative strategy, we fused eGFP::LOV to endogenous PAR-6, which localizes to the anterior cortex of the zygote (Fig. 3a). Following global light exposure, recruitment of RGS-7::ePDZ to PAR-6::LOV enhanced the peak velocities of both spindle poles, but most significantly the movement of the anterior pole (anterior +25% and posterior +14%; Fig. 3b, c). Thus, cortical recruitment of RGS-7 acutely increases pulling forces, in agreement with its proposed function as a GAP that promotes Gα∙GDP–GPR-1/2 interaction. In contrast, cortical enrichment of RIC-8 did not significantly enhance pulling forces (Fig. 3e). Thus, in agreement with the *ric-8^lox^; rgs-7^lox^* synergistic phenotype, our optogenetic localization experiments support a model in which RIC-8 and RGS-7 promote Gα function at different levels. While RGS-7 probably functions as a GAP, our data are in line with RIC-8 acting in vivo as a Gα chaperone, rather than a GEF, thus indirectly promoting force generation.

**Figure 3.**
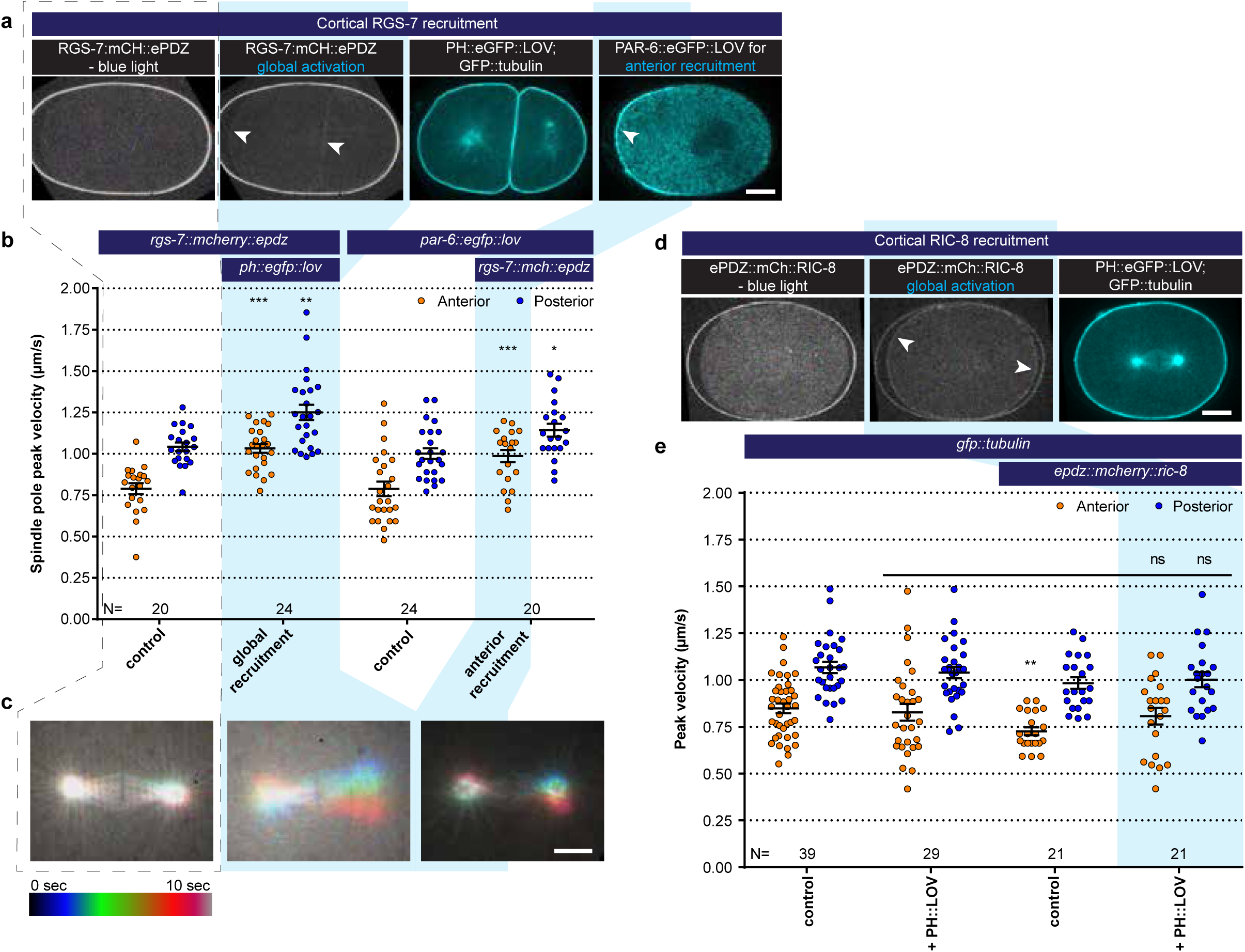
Light-controlled localizaon of endogenous Gα regulatory proteins in the *C. elegans* embryo. **(a)** Light-controlled localizaon of endogenous RGS-7 to membrane PH::LOV (arrowheads, note of the auto fluorescent eggshell in the mCherry channel). Most right panel: anterior localizaon of PAR-6::eGFP::LOV. Scale bar: 10 µm **(b)** Spindle severing experiments aer light-induced corcal localizaon of RGS-7 (blue fields). Controls are *rgs-7::mcherry::epdz* and *par-6::egfp::lov*. Experimental condions: combinaon with *ph::egfp::lov* and *rgs-7::mcherry::epdz*. Blue light acvaon was global and connuous. Error bars: s.e.m. Welch’s Student’s t-test; * P < 0.05, ** P < 0.01, *** P < 0.001. **(c)** Maximum projections of spindle movements for 10 seconds using a temporal color coding scheme to visualize spindle movement. A staonary spindle produces a white maximum projecon, whereas a mobile spindle leaves a colored trace. Scale bar: 5 µm. **(d)** Light-controlled localizaon of endogenous RIC-8 to membrane PH::eGFP::LOV (arrowheads, note of the auto fluorescent eggshell in the mCherry channel). Scale bar: 10 µm **(e)** Spindle severing experiments aer light-induced corcal localizaon of RIC-8 (blue fields). Controls are *gfp::tubulin* and *epdz::mcherry::ric-8*. Experimental condions: combinaon with *ph::egfp::lov*. Blue light acvaon was global and connuous. Error bars: s.e.m. Welch’s Student’s t-test; ns P > 0.05. Blue fields indicate condions where an ePDZ-LOV interacon was induced. See Supplementary Table 1 for detailed genotypes. Anterior is to the le in microscopy images.

To directly address whether Gα∙GTP might contribute to spindle positioning or if Gα∙GDP serves merely as a membrane anchor, we aimed to reconstruct a cortical force generator in the absence of Gα (Fig. 4a). We obtained optogenetic control over the membrane localization of GPR-1 by combining endogenously labeled *epdz::mcherry::gpr-1* (ePDZ::GPR-1) with knockout of *gpr-2*, expression of PH::LOV and Gα RNAi (Fig. 4b, c and Supplementary Video 8, 9). Live imaging and immunohistochemistry confirmed light-induced cortical recruitment of ePDZ::GPR-1 and consequently LIN-5 (Fig. 4b, Fig. S3a). Spindle pulling forces appeared reduced following the tagging of *gpr-1* and knockout of *gpr-2* (anterior −40% and posterior −26%) (Fig. S3b). However, light-induced ePDZ::GPR-1 recruitment increased spindle pole movements (anterior +56% and posterior +10%) (Fig. 4d, f). Moreover, membrane-localized ePDZ::GPR-1 sustained force generation in the absence of Gα (anterior +195% and posterior +232%), indicating that Gα is dispensable for cortical pulling force generation. Recruitment of ePDZ::GPR-1 restored spindle pole movements to a similar degree in *Gα(RNAi)* and *Gα(RNAi); ric-8(RNAi)* embryos (Fig. 4d). Thus, cortical pulling forces can be generated when Gα membrane anchor function is replaced by PH::LOV, and most likely in the absence of Gα_i/o_∙GTP. We conclude that Gα functions as a membrane anchor and that Gα∙GTP does not perform an essential function in pulling force generation.

**Figure 4.**
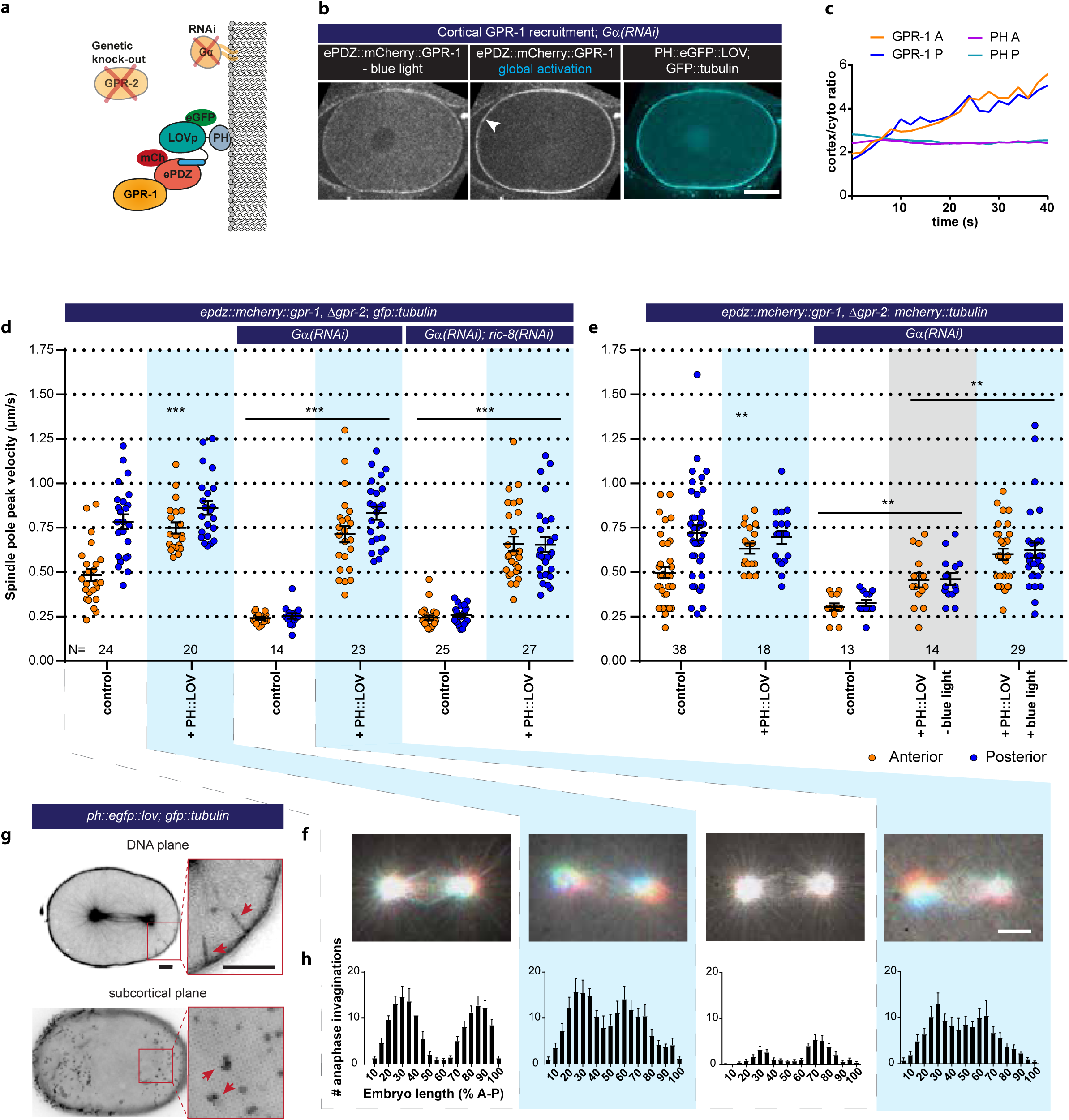
Light-inducible GPR-1 recruitment to the cortex rescues pulling force generaon in the absence of Gα. **(a)** Cartoon model illustrang the experiment that localizes GPR-1 directly to the membrane, bypassing the wild type membrane anchor Gα which is inacvated by RNAi. **(b)** Spinning disk confocal images of light-controlled corcal GPR-1 recruitment independent of the wild type anchor Gα (arrowheads; note the auto fluorescent eggshell in the mCherry channel). Scale bar: 10 µm. **(c)** Quanficaon of corcal GPR-1 recruitment during connuous acvaon of the ePDZ-LOV interacon, represented as the rao of corcal/cytoplasmic signal. Also see Supplementary Movie 8, 9. **(d)** Spindle severing experiments in combinaon with corcal recruitment of endogenous GPR-1. Control is *epdz::mch::gpr-1, Δgpr-2; gfp::tubulin*. Experimental condions: combinaons with *ph::egfp::lov, Gα(RNAi), and Gα(RNAi); ric-8(RNAi)*. Blue light acvaon was global and connuous. Error bars: s.e.m. Welch’s Student’s t-test and Mann Whitney U test; ** P < 0.01, *** P < 0.001. **(d)** Spindle severing experiments in combinaon with corcal recruitment of endogenous GPR-1. Control is *epdz::mch::gpr-1, Δgpr-2; mcherry::tubulin*. Experimental condions: combinaons with *ph::egfp::lov, G α(RNAi)*, and the absence of blue light (grey field). Blue light acvaon was global and connuous. Error bars: s.e.m. Welch’s Student’s t-test and Mann Whitney U test; ** P < 0.01, *** P < 0.001. **(e)** Maximum projecons of spindle movements for 10 seconds using a temporal color coding scheme to visualize spindle movement as in Fig. 3c. **(f)** Plasma membrane invaginaons as result of corcal pulling forces are visible as lines in the DNA plane and dots in the subcorcal plane (red arrows). Scale bar: 5 µm. **(g)** Distribuon of anaphase membrane invaginaons ploed along anterior-posterior embryo length. Condions were the same as for the connected experiments in e and f, except for the control which was *ph::eg-fp::lov; gfp::tubulin* and not *epdz::mch::gpr-1, Δgpr-2*. Scale bar: 5 µm. Blue fields indicate condions where an ePDZ-LOV interacon was induced. See Supplementary Table 1 for detailed genotypes. Anterior is to the le in microscopy images.

Light-controlled heterodimerization exhibits a certain level of dark state activity (Hallett et al., 2016). We performed spindle severing experiments in the absence of blue light to confirm the light-specificity of recruitment. Replacement of *gfp::tubulin* with an *mcherry::tubulin* transgene allowed for tracking of the spindle in the absence of blue light and consequently LOV activation. We observed that the scattering of UV-light (355 nm) during midzone ablation also uncages the LOV domain. Nevertheless, the presence of blue light resulted in substantially elevated spindle pulling forces when compared to dark state experiments (anterior +33% and posterior +35%) (Fig. 4e). We conclude that the observed spindle pole movements are light-dependent and the specific result of inducible cortical recruitment of GPR-1.

Considering that spindle poles moved in three dimensions after recruitment of GPR-1, measuring peak velocities after midzone severing by tracking the poles in two dimensions likely underestimated the resulting pulling forces. Therefore, we utilized an additional read-out of cortical pulling forces. Cortical pulling events cause invaginations of the plasma membrane (Redemann et al., 2010), which are visible as dots in the sub-cortical plane (Fig. 4g and Supplementary Video 10, 11). Control PH::LOV embryos showed on average 138 membrane invaginations during anaphase in an area covering approximately 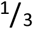 of the cell surface (Fig. S3d). When plotted along the anterior-posterior axis, the distribution of these invaginations reflected the three described cortical domains: anterior, posterior, and a posterior lateral LET-99 region at ±60% embryo length (Rose and Gönczy, 2014) (Fig. 4h). In this cortical domain, LET-99 antagonizes GPR-1/2 localization and thereby pulling force generation (Krueger et al., 2010). Cortical GPR-1 recruitment resulted in a total number of 174 (+25% compared to *ph::lov* control) invaginations in the presence and 122 (+249% compared to *Gα(RNAi)*) invaginations in the absence of Gα (Fig. S3c, d). Interestingly, the absence of invaginations around 60% embryo length was lost following ePDZ::GPR-1 anchoring by PH::LOV. Instead, we observed a milder dip at 50% embryo length. The remaining peaks likely represent the cortical regions closest to the spindle poles, as these sites contact the highest numbers of astral microtubules. Taken together, Gα is not essential for force generation, but the characteristic distribution of force generating events is likely regulated in part at the Gα level.

Gα–GPR-1/2–LIN-5 has been suggested to function as a dynein anchor (Kotak et al., 2012; Pietro et al., 2016), but this has not been tested directly. Our optogenetic approach allows replacing the entire complex by PH::LOV, and examining whether the complex strictly acts as an anchor, or whether individual components have additional functions (Fig. 5a). To directly recruit dynein to the cortex, we generated an *epdz::mcherry* knock-in allele of *dhc-1* (dynein heavy chain). While homozygous *epdz::mcherry::dhc-1* (ePDZ::DHC-1) was viable, its combination with *ph::egfp::lov* was lethal. This effect was also observed for an ePDZ::GFP fusion of DHC-1 in the presence of PH::LOV, but not in the absence of PH::LOV or for mCherry::DHC-1 without the ePDZ domain. Therefore, we attributed the lethality to ePDZ-LOV dark state interactions that disturb essential dynein functions. We circumvented this effect by using *epdz::mcherry::dhc-1* in combination with a wild-type allele (*epdz::mcherry::dhc-1/+*) to control dynein localization (Fig. 5b, c and Supplementary Video 12, 13). We found that induced ePDZ::DHC-1 cortical recruitment in the presence of the wild type complex slightly (but not significantly) increased spindle pole movements (anterior +10% and posterior +5%, 162 membrane invaginations: +17%; Fig. 5d-f). Notably, cortical ePDZ::DHC-1 recruitment in the absence of a wild type complex (*gpr-1/2(RNAi)* or *Gα(RNAi)*) did not result in substantial pulling force generation or spindle movements (Fig. 4e, Fig. S4b, c). Because direct cortical dynein anchoring does not support force generation, it is likely that the LIN-5 complex performs essential functions beyond providing a structural dynein anchor.

**Figure 5.**
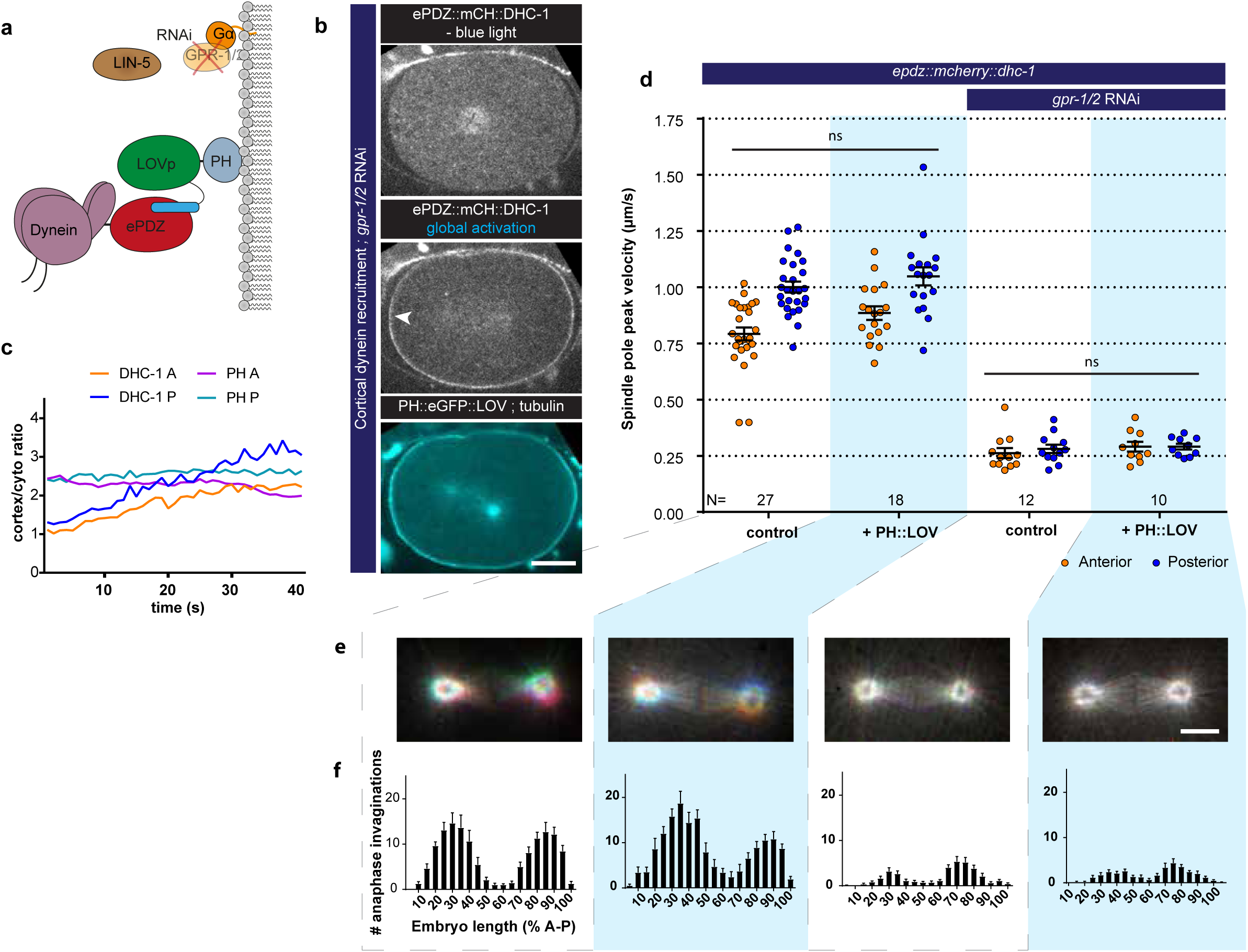
Direct corcal anchoring of dynein (DHC-1) is insufficient for corcal pulling force generaon. **(a)** Cartoon model illustrang the experiment where dynein is recruited directly to the cortex. The wild type force generator is inacvated by RNAi. **(b)** Spinning disk confocal images showing light-controlled recruitment of dynein to the cortex (arrowheads, note the auto fluorescent eggshell in the mCherry channel). Scale bar: 10 µm. **(c)** Quanficaon of corcal dynein recruitment during connuous acvaon of PH::LOV with blue light. (d) Spindle severing experiments with corcal dynein recruitment. Control is *epdz::mch::dhc-1; gfp::tubulin*. Experimental condions: combination with *ph::egfp::lov* and *gpr-1/2(RNAi)*. Blue light acvaon was global and connuous. Error bars: s.e.m. Welch’s Student’s t-test and Mann Whitney U test; ns P > 0.05. (e) Maximum projecons of spindle movements for 10 seconds using a temporal color coding scheme to visualize spindle movement as in Figure 3c. Scale bar: 5 µm. (f) Distribuon of anaphase membrane invaginaons ploed along anterior-posterior embryo length. Condions were the same as for the connected experiments in d and e, except for the control which was *ph::egfp::lov; gfp::tubulin* and not *epdz::mch::dhc-1*. Blue fields indicate condions where an ePDZ-LOV interacon was induced. See Supplementary Table 1 for detailed genotypes. Anterior is to the le in microscopy images.

*In vitro* reconstitution studies established that homodimerizing adapters containing extended coiled-coil domains are critical for dynein activity (McKenney et al., 2014; Schlager et al., 2014). LIN-5 and its homologs NuMA and Mud are predicted to contain a long coiled-coil domain, to homodimerize, and to interact with dynein (Kotak et al., 2012; Lorson et al., 2000; Merdes et al., 1996). To investigate if LIN-5 can activate dynein-dependent force generation, we recruited endogenous LIN-5 to the cortex (Fig. 6a-c and Supplementary Video 14, 15). Spindle severing experiments and invagination counting revealed that cortical LIN-5 recruitment greatly increased spindle pulling forces in otherwise wild type embryos (anterior +131% and posterior +68%, 557 invaginations: +303%) (Fig. 6d, e and Fig. S5b-d). *gpr-1/2*(*RNAi)* embryos also showed strong dynein-dependent forces after cortical recruitment of LIN-5 (anterior +183% and posterior +244%, 429 invaginations: +1488%). In fact, cortical LIN-5 recruitment generated extreme premature pulling forces (Supplementary Video 16) that could result in separation of centrosomes and their associated pronuclei even before formation of a bipolar spindle (Supplementary Video 17). Therefore, embryos were kept in the absence of blue light until mitotic metaphase. Subsequent blue light exposure induced cortical LIN-5 recruitment within seconds, and the spindles showed excessive movements in all three dimensions well before cortical LIN-5 reached peak levels (Fig. 6c, e). Therefore, the number of membrane invaginations in anaphase probably reflects the pulling forces more accurately than the average peak velocities of the poles (Fig. 6f and Fig. S5b, c). These results identify LIN-5 as a strong activator of dynein in the generation of cortical pulling forces.

**Figure 6.**
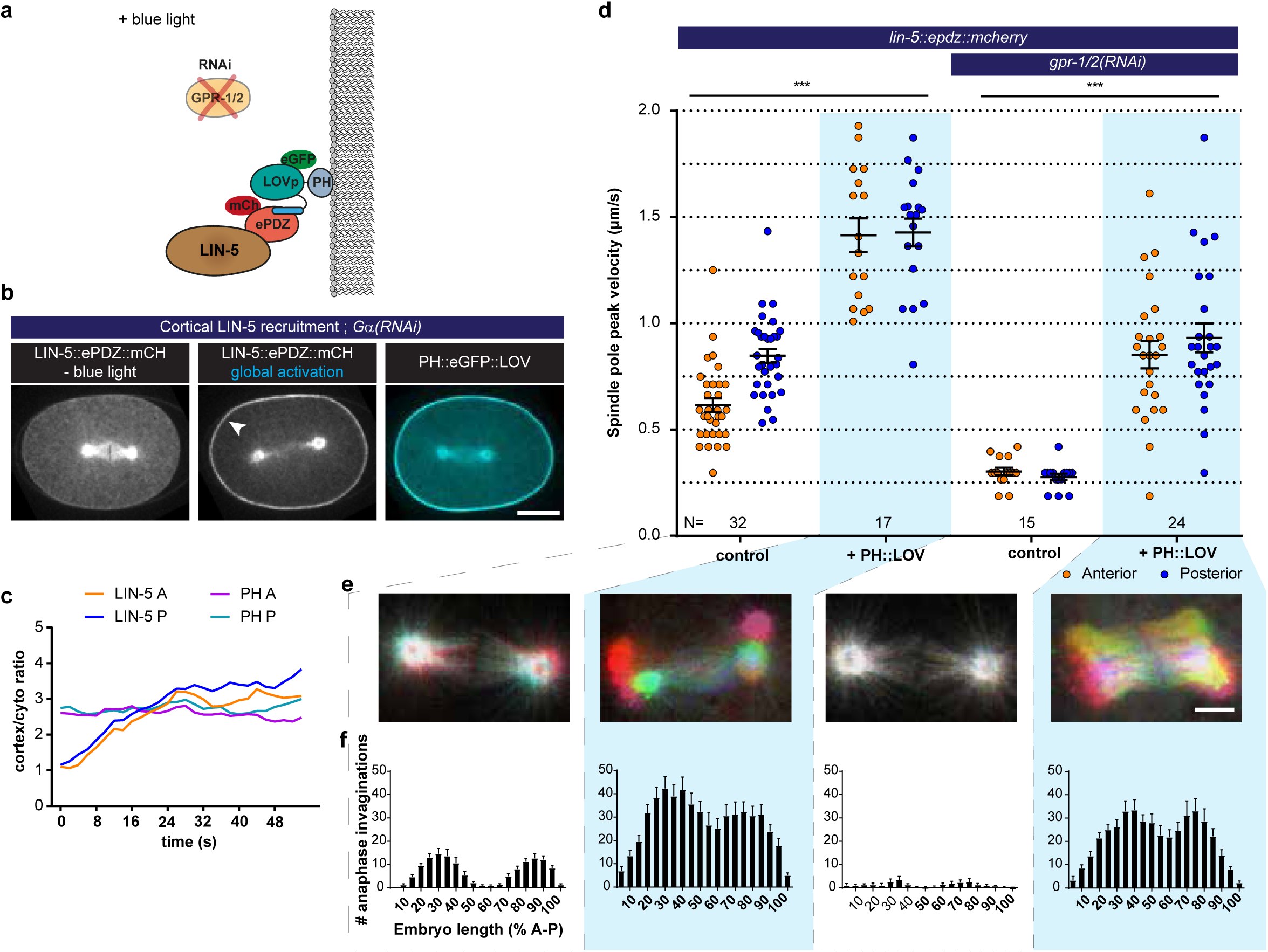
LIN-5 is a strong and essenal acvator of dynein-dependent corcal pulling forces. **(a)** Cartoon model illustrang the experiment in which LIN-5 is recruited to the cortex independently of the wild type Gα-GPR-1/2 anchor. **(b)** Spinning disk confocal images showing light-controlled recruitment of endogenous LIN-5 in the absence of Gα (arrow head). See also Supplementary Movie 14, 15. **(c)** Corcal LIN-5 recruitment during connuous acvaon of the ePDZ-LOV interacon, represented as the rao of corcal/cytoplasmic signal. Scale bar: 10 µm. **(d)** Spindle severing experiments in combinaon with corcal recruitment of endogenous LIN-5. Control was *lin-5::epdz::mcherry; gfp::tubulin*. Experimental condions: combinaons with *ph::egfp::lov* and *gpr-1/2(RNAi)*. Error bars: s.e.m. Welch’s Student’s t-test and Mann Whitney U test; *** P < 0.001. (e) Maximum projecons of spindle movements for 10 seconds using a temporal color coding scheme to visualize spindle movement as in Fig. 3c. Scale bar: 5 µm. (f) Anaphase membrane invaginaons ploed along anterior-posterior embryo length. Condions were the same as for the connected experiments in d and e, except for the control which was *ph::egfp::lov; gfp::tubulin* and not *lin-5::ep-dz::mch*. Blue fields indicate condions where an ePDZ-LOV interacon was induced. See Supplementary Table 1 for detailed genotypes. Anterior is to the le in microscopy images.

Next, we examined whether we could deploy cortical LIN-5 to manipulate the spindle position and outcome of cell division by local illumination with blue light. In the normal P0 cell, the spindle becomes positioned posteriorly and cell cleavage creates a larger anterior blastomere (AB) and smaller posterior blastomere (P1). Local recruitment of LIN-5 to the anterior cortex caused the P0 spindle to position anteriorly and inverted the AB:P1 asymmetry (Fig. 7a and Supplementary Video 18). In addition, recruiting LIN-5 laterally induced a completely perpendicular spindle position (Fig. 7b and Supplementary Video 19). While this triggered some furrowing at the anterior and posterior cell poles, the spindle switched back to an anterior-posterior orientation during cytokinesis, possibly resulting from geometric constraints. We therefore switched to two-cell embryos with the relatively round AB and P1 blastomeres. In two-cell *gpr-1/2(RNAi)* embryos, the spindle fails to rotate in P1, resulting in a transverse spindle orientation in both blastomeres (Srinivasan et al., 2003). Importantly, local recruitment of LIN-5 to the membranes between AB and P1 promoted anterior-posterior spindle orientations in both blastomeres of *gpr-1/2(RNAi)* embryos (Fig. 7c and Supplementary Video 20, 21). These spindles maintained their anterior-posterior orientation throughout mitosis and induced cleavage furrows that reproducibly followed the spindle positions. These experiments underline the determining role of LIN-5-dependent cortical pulling in spindle orientation and cell cleavage determination.

**Figure 7.**
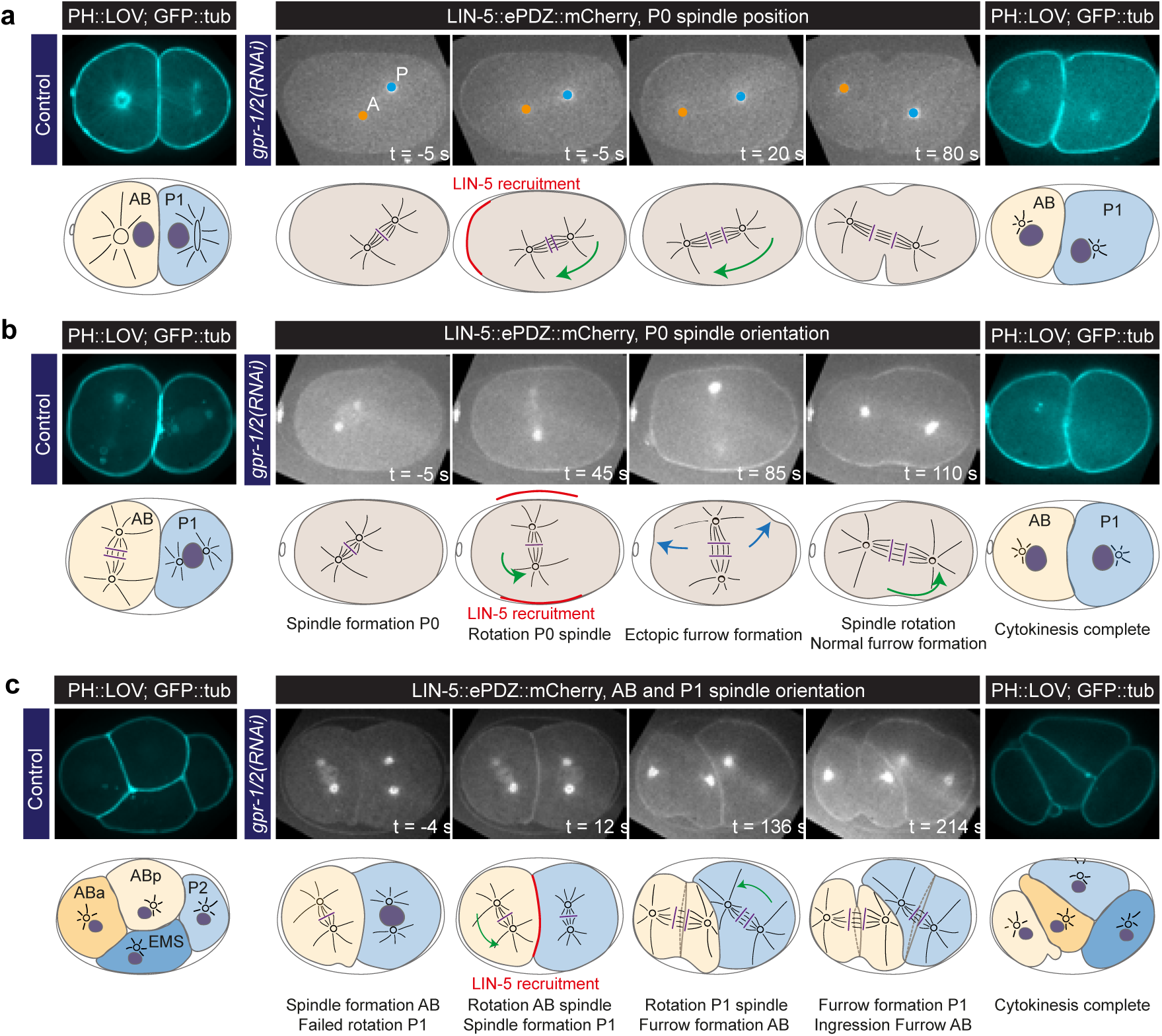
Experimentally induced spindle posioning by controlled localizaon of endogenous LIN-5. **(a)** Selected me points of Supplementary Movie 18 showing induced anterior displacement of the P0 spindle upon local corcal recruitment of LIN-5. Images are annotated with centrosome posions shown as circles (orange, anterior pole; blue, posterior pole). **(b)** Selected me points of Supplementary Movie 19 showing induced transverse P0 spindle orientaon upon local corcal recruitment of LIN-5. Blue arrows, ectopic furrowing. **(c)** Selected me points of Supplementary Movie 20 showing induced AB and P1 spindle rotaon upon local corcal recruitment of LIN-5. In a, b, and c panels 2–5 show LIN-5::eP-DZ::mCherry fluorescence, panel 1 and 5 show PH::LOV and GFP::tubulin. Cartoons accompanying images illustrate key events. Red, local LIN-5 recruitment. Green arrows, spindle movements. lemost panels show control 2- and 4-cell embryos labeled with PH::eGFP::LOV and GFP::Tubulin. See Supplementary Table 1 for detailed genotypes. Anterior is to the left.

Our observations fit with and expand on conclusions from studies of LIN-5-related Mud and NuMA, which when tethered to the cortex contributes to spindle positioning (Kotak et al., 2012; Ségalen et al., 2010). Notably, these previous studies considered NuMA to be just a dynein anchor, and were performed in cells that express Gα, GPR-related Pins and LGN, and presumably RIC-8A proteins. We observed similarly high pulling forces upon cortical LIN-5 recruitment in *gpr-1/2(RNAi)* and *Gα(RNAi)* embryos (Fig. S5c, d). Thus, Gα and GPR-1/2 are not per se required for force generation. In addition, we observed that cortical recruitment of LIN-5 localized dynein and vice versa (Fig. S4a, S5a). However, only LIN-5-mediated cortical anchoring of dynein through LIN-5 resulted in pulling forces. Thus, membrane attachment of LIN-5 appears both essential and sufficient for dynein-dependent cortical pulling force generation.

## DISCUSSION

Recent advances in CRISPR/Cas9-mediated genome engineering (Waaijers and Boxem, 2014) and optogenetics (Johnson and Toettcher, 2018) hold far-reaching potential for cell- and developmental biology. We combined these strategies to systematically control the localization of endogenous proteins in the *C. elegans* early embryo by light-induced ePDZ-LOV heterodimerization, to determine their individual contributions in spindle positioning. Our quantitative analyses classified LIN-5, but not Gα, RIC-8 and GPR-1/2, as essential for dynein-dependent pulling force generation. Since only LIN-5 is strictly required for cortical pulling force generation, the question arises why a tripartite dynein anchor is conserved from worm to man. In yeast, dynein is localized by the single-component cortical anchor Num1, a coiled-coil domain protein with a PH-domain for membrane localization (Ananthanarayanan, 2016). Our ectopic ePDZ-LOV heterodimerization experiments show that membrane-tethered LIN-5 could suffice as a dynein anchor and activator, and that local regulation is needed to rotate and migrate the spindle. Conceivably, the trimeric dynein anchor/adaptor evolved in metazoans to augment context-specific regulation and reduce stochastic activation of spindle pulling forces. We propose that Gα–GPR-1/2 provides a regulatable membrane anchor, while membrane-bound LIN-5 acts as an obligate adapter and activator of cytoplasmic dynein at the cell cortex, which possibly links dynein and the dynactin complex.

Which factors may normally control the Gα–GPR-1/2 membrane anchor? Our double knock-out experiments and optogenetic localization studies support that RIC-8 and RGS-7 regulate Gα anchor function at different levels. RIC-8 was initially characterized as a GEF for Gα_i/o_∙GDP (Tall et al., 2003) and Gα∙GDP-LGN-NuMA (Tall and Gilman, 2005). To explain how a GEF could promote spindle pulling forces, it has been proposed that Gα∙GDP/GTP cycling is essential for Gα∙GDP-GPR-1/2 association. However, experiments in *Drosophila* (David et al., 2005; Hampoelz et al., 2005; Wang et al., 2005) and mammals (Gabay et al., 2011) suggested a Gα chaperone function for RIC-8, while observations in *C. elegans* supported dual activities as a GEF for GOA-1^Gαo^ (Afshar et al., 2004), and a chaperone for the membrane localization of GPA-16^Gαi^ (Afshar et al., 2005). We found no evidence to support direct RIC-8 function in spindle positioning in vivo and favor a model in which RIC-8 indirectly affects force generation by promoting the properly folded membrane-anchored Gα_i/o_ conformation, possibly as chaperone. The fact that Gα can be replaced with a PH-membrane anchor dismisses a general requirement for Gα∙GTP in pulling force generation. However, we noticed that Gα contributes to the normal distribution of pulling forces: direct cortical anchoring of GPR-1 or LIN-5 coincided with increased membrane invaginations in the region occupied by LET-99. This indicates that LET-99 normally antagonizes Gα∙GDP availability or Gα∙GDP-GPR-1/2 interaction.

Despite the observed replaceability of Gα, Gα∙GTP has been reported to affect the spindle orientation in specific tissues (Katanaev et al., 2005; Schaefer et al., 2001; Yoshiura et al., 2012). In *Drosophila* neuroblasts and sensory organ precursor cells, canonical G-protein signaling probably is used to align cellular polarity with tissue polarity (Katanaev et al., 2005; Yoshiura et al., 2012).

For our in vivo dissection of spindle positioning, we developed and applied methods for germline-specific gene knockout, tagging of endogenous proteins, reliable expression of foreign sequences in the germline, and light-inducible protein heterodimerization. These methods further expand the molecular biology toolbox for in vivo studies and can be broadly applied to other biological processes. Of particular interest is the acquired possibility to experimentally control the position of the spindle, for instance for future studies aimed at deciphering how the spindle determines the plane of cell cleavage.

## MATERIALS and METHODS

### *C. elegans* strains and maintenance

The names and associated genotypes of *C. elegans* strains used in this study are included in supplementary table 1. Animals were maintained at either 15 or 20 ˚C as described previously (Brenner, 1974). Strains expressing both ePDZ and LOV protein motifs were regarded as light-sensitive and thus cultured in the dark. Animals were kept on plates that contained nematode growth medium (NGM) that had been seeded with OP50 *Escherichia coli* bacteria.

### Molecular cloning

DNA vector-based repair templates to be used for CRISPR/Cas-9-mediated genome editing were designed in A plasmid Editor (M. Wayne Davis) to include 500–1500 bp homology arms. These and all other sequences used were generated using either purified *C. elegans* genomic DNA or pre-existing vectors via PCR amplification using Q5 Hot Start High-Fidelity DNA Polymerase (New England Biolabs). A list of all cloning, repair template and genotyping primers (Integrated DNA technologies) and DNA templates used has been included in supplementary table 2. PCR fragments were gel purified (Qiagen), their concentrations measured using a BioPhotometer D30 (Eppendorf) and then ligated into pBSK by Gibson assembly (New England Biolabs). gRNA vectors were generated by annealing of antisense oligonucleotide pairs and subsequent ligation into BbsI-linearized pJJR50 or BsaI-linearized pMB70 using T4 ligase (New England Biolabs). All DNA vectors used for genome editing were transformed into DH5α competent cells and subsequently purified by midiprep (Qiagen).

### Design of germline-optimized coding sequences

Custom Perl scripts were written to design germline-optimized coding sequences according to the algorithm described in the legend of Supplementary Fig. 2. After designing each coding sequence, we inserted either 1) normal synthetic introns with the sequence gtaagttt(n36)ttttcag, where n36 is a 36 bp random DNA sequence with 30% GC content; or 2) PATC introns (Frøkjær-jensen et al., 2016). Our design algorithm is accessible via a web interface at http://104.131.81.59/, and the source code can be found at https://github.com/dannyhmg/germline. Germline-optimized sequences were synthesized as gBlocks (Integrated DNA Technologies) and single-copy transgenes were generated using standard methods (Frøkjær-Jensen et al., 2012). Please refer to supplementary table 2 for detailed sequence features of each transgene.

### Design of inducible germline-specific knockout

*loxP* and *loxN* sequences were integrated in the endogenous loci of essential genes (see CRISPR/Cas9-mediated genome editing section for details). For the FLP recombinase, the hyperactive FLP G5D variant (Schwartz and Jorgensen, 2016) was used (pMLS262; Addgene #73718). We used a long version of the *pie-1* promoter including enhancer for germline-specific expression (pAZ132, a kind gift from A. A. Hyman). To initiate the recombination cascade, the germline-specific FLP was injected in P0 mothers with the following protocol to favor germline expression (Personal communication Oliver Hobert, February 21, 2013); linearized FLP construct (2 ng/µl), PvuII digested *E. coli* genomic DNA (150 ng/µl), co-injection marker *Pmyo-2::tdtomato* (2 ng/µl). Transgenic F1 animals were singled and allowed to lay eggs for at least 24 hrs. From F1 with 100% embryonic lethal broods (*ric-8* and *rgs-7* are essential for embryogenesis), early embryos were isolated and used for spindle severing experiments.

### CRISPR/Cas9-mediated genome editing

Either the wild type N2 or SV1818 (*pha-1*(*e2123*ts) 4x outcrossed) *C. elegans* genetic background was used for the generation of CRISPR/Cas-9 alleles. Injection mixes with a total volume of 50 µl were prepared in MilliQ H_2_O and contained a combination of 50 ng/µl *Peft-3-p::cas9* (Addgene ID #46168 (Martin and Weiss, 2015) or 60 ng/µl pJW1285 (61252, Addgene; (Dickinson et al., 2013)), 50–100 ng/µl *u6::sgRNA* (targeting genomic sequences listed in supplementary table 2), 50 ng/µl of (PAGE-purified oligonucleotide) repair template and 2.5 ng/µl of the co-injection pharyngeal marker *myo-2p::tdtomato*, and spun down in a microcentrifuge (Eppendorf) for at least 10 minutes at 13,000 RPM prior to use. Young adult hermaphrodites were injected in the germline using an inverted micro-injection setup (Eppendorf). After injection, animals were singled and grown at 15 or 20 ˚C. F1 animals were then picked to a total of at least 96, and grown with two or three animals per plate for 7–8 days at 20 ˚C until freshly starved. Half a plate containing F2 and F3 animals was then washed off with M9 medium supplemented with 0.05% Tween-20, and subsequently lysed to extract genomic DNA. Some knock-ins were obtained using co-CRISPR selection: rescue of *pha-1*(*e2123*ts) (Ward, 2014), generation of visible *unc-22* (Kim et al., 2014) or *dpy-10* (Paix et al., 2015) phenotypes, or integration of an self-excisable cassette carrying a visible marker (Dickinson et al., 2015). Genotyping was carried out by PCR amplification with OneTaq polymerase (New England Biolabs) of genome sequences using primers annealing in the inserted sequence and a genomic region not included in the repair template. Confirmed alleles were subsequently sequenced (Macrogen Europe).

### Spinning disk microscopy

Prior to live imaging, embryos were dissected from adult hermaphrodites onto coverslips (Menzel-Gläser) in 0.8x egg salts buffer (94 mM NaCl, 32 mM KCl, 2.7 mM CaCl2, 2.7 mM MgCl2, 4 mM HEPES, pH 7.5; (Tagawa et al., 2001) or M9, and mounted on 4% agarose pads. Spinning disk imaging of embryos was performed using a Nikon Eclipse Ti with Perfect Focus System, Yokogawa CSU-X1-A1 spinning disk confocal head, Plan Apo VC 60x N.A. 1.40 oil and S Fluor 100x N.A. 0.5–1.3 (at 1.3, used for UV-laser photo-ablation) objectives, Photometrics Evolve 512 EMCCD camera, DV2 two-channel beam-splitter for simultaneous dual-color imaging, Cobolt Calypso 491 nm (100 mW), Cobolt Jive 561 nm (100 mW) and Teem Photonics 355 nm Q-switched pulsed laser controlled with the ILas system (Roper Scientific France/PICT-IBiSA, Institut Curie, used for photo-ablation), ET-GFP (49002), ET-mCherry (49008) and ET-GFPmCherry (49022) filters, ASI motorized stage MS-2000-XYZ with Piezo Top Plate, and Sutter LB10-3 filter wheel. The microscope was operated using MetaMorph 7.7 software and situated in a temperature-controlled room (20 ˚C). The temperature of the stage and objective was controlled at 25 ˚C with a Tokai Hit INUBG2E-ZILCS Stage Top Incubator during experiments. Images were acquired in either streaming mode with 250 or 500 ms exposure, or time-lapse mode with 250, 500 or 1500 ms exposure and 2 or 5 second intervals. Laser power and exposure times were kept constant within experiments. For the quantification of membrane invaginations embryos were imaged by 250 ms exposure stream acquisition starting in the DNA plane at anaphase onset, as judged by GFP::Tubulin signal. During anaphase, the spinning disk imaging plane was moved as close to the membrane as possible while keeping the cytosol discernable from the membrane signal. Acquisitions were terminated at early telophase, as judged by the PH::eGFP::LOV signal. For experiments involving balanced *epdz::mcherry::dhc-1/+*, each animal was confirmed to be positive for *epdz::mcherry::dhc-1* by fluorescence before the experiment. Images acquired by spinning disk microscopy were rotated, cropped, annotated, provided with scale bars, and processed further by linear adjustment of brightness and contrast using ImageJ and FIJI. Fluorophores used in this study include (e)GFP, mCherry, Alexa-488 and Alexa-568.

### RNA-mediated interference (RNAi)

For immunohistochemistry experiments L4 hermaphrodites were grown on RNAi plates seeded with HT115 *Escherichia coli* bacteria strains generating double-stranded RNA (dsRNA) targeting genes of interest (*goa-1, gpa-16, gpr-1*) for 48 hours at 15 ˚C prior to fixation (Timmons and Fire, 1998). For all other gene knock-down experiments, young adult hermaphrodites were injected with dsRNA targeting genes of interest (*goa-1, gpa-16, gpr-1, ric-8, rgs-7*) and grown for 48 hours at 15 ˚C (Fire et al., 1998) prior to experiments. To generate dsRNA, coding regions of genes of interest were PCR amplified using Q5 Hot Start High-Fidelity DNA Polymerase (New England Biolabs). These PCR products were used as templates for *in vitro* dsRNA synthesis (MEGAscript T7 transcription kit, ThermoFisher Scientific). dsRNA was diluted 5x in DEPC H_2_O prior to micro-injection. ORF clones from the Vidal and Ahringer RNAi libraries were used (Kamath et al., 2003; Rual et al., 2004).

### Spindle severing assays

Mitotic spindle severing was performed in essence as described (Grill et al., 2001; Portegijs et al., 2016). One-cell embryos expressing GFP-or mCherry-labeled tubulin were imaged during mitosis using the spinning disk microscope setup described above, equipped with a Teem Photonics 355 nm Q-switched pulsed laser controlled with the ILas system (Roper Scientific France/PICT-IBiSA, Institut Curie). At anaphase onset, as judged by spindle morphology and mobility, spindles were severed as shown in Figure 1c and Supplementary Movie 2. Centrosome displacement was recorded by 500 ms exposure streaming acquisition, and peak velocities were subsequently extrapolated using the FIJI TrackMate plugin.

### Dark state spinning disk microscopy

Dark state experiments were performed on the spinning disk setup described above. For local photoactivation of LOV2 in *C. elegans* embryos, light was applied in a region of variable size depending on each individual experiment using a 491 nm laser controlled with the ILas system (Roper Scientific France/ PICT-IBiSA, Institut Curie). Due to high sensitivity of LOV2 to blue light and variations in laser power, embryos of strain SV2061 (expressing diffuse ePDZ::mCherry and PH::eGFP::LOV) were used to calibrate the amount of laser power required for local activation of LOV2 prior to experiments. During both global and local photoactivation assays and dark state spindle severing experiments embryos were kept away from blue light as much as practically feasible. To this end, aluminum foil was used to cover the microscope setup, and optical filters were inserted in the light path to remove LOV2-activating wavelengths from the transmitted light used to locate embryos on slides. Prior to experimental use of embryos, unintended premature cortical recruitment of ePDZ-mCherry or ePDZ-mCherry-LIN-5 was assessed by observation of mCherry localization patterns.

### Antibodies and immunocytochemistry

For immunostaining of *C. elegans* embryos, embryos were dissected from adults in 10 µl MilliQ H_2_O on slides coated with poly-L-lysine. Samples were then freeze-cracked and fixed in methanol for 5 min. at −20 ˚C and subsequently in acetone for 5 min. at −20 ˚C. Embryos were then rehydrated in phosphate buffered saline + 0.05% Tween-20 (PBST), blocked for 1 hour at 4 ˚C in PBST + 1% bovine serum albumin and 1% goat serum (Sigma-Aldrich), and then incubated at room temperature with primary antibodies for 1 hour and then with secondary antibodies for 45 min., both in blocking solution, with four 10 minute washes in PBST following each antibody mix. Finally, embryos were embedded in ProLong Gold Antifade with DAPI. Primary antibodies used were mouse anti-LIN-5 (1:10, (Lorson et al., 2000)) and rabbit anti-DHC-1 (1:100, (Gönczy et al., 1999); a kind gift from P. Gönczy). Secondary antibodies used were goat anti-rabbit Alexa-488, goat anti-rabbit Alexa-568, goat anti-mouse Alexa-488 and goat anti-mouse Alexa-568 (Invitrogen), all at 1:500 dilution. Imaging of immunolabeled embryos was performed on the spinning disk setup described above.

### Data analysis

All quantitative spinning disk image analyses were performed in either ImageJ or FIJI. For quantification of membrane invaginations, movies were limited to the 200 frames (50 seconds) preceding the onset of telophase. Images were then cropped to include the outer limits of the PH::eGFP signal. Transient cortical dots were tracked manually using the MTrackJ ImageJ plugin. To yield the distribution of invaginations on the length axis of the visible embryo cortex, recorded x coordinates were incremented into groups of 5% embryo length each. To quantify the cortical recruitment and dynamics of ePDZ::mCherry, ePDZ::mCherry::GPR-1, LIN-5::ePDZ::mCherry and ePDZ::mCherry::DHC-1 by PH::eGFP::LOV, multiple 20 px wide linescans were drawn perpendicular to the membrane per analyzed embryo. An intensity profile was plotted per linescan at each acquired time point, from each of which an average of the maximum 3 pixel values was extracted to yield the peak intensity values at the membrane. Each intensity measurement was first corrected for background noise with a value measured outside of the embryo in a 50×50 px region of interest, and cortex to cytoplasm intensity ratios were calculated using average cytoplasmic intensity measurements in a 50×50 or 29×23 px region of interest at all timepoints analyzed. Fluorescence intensity measurements as measure for *Cre(FLPon)* activation (Supplementary Fig. 1e) were taken as total embryo average intensity minus background signal using ImageJ measurement tool. The half time of ePDZ-LOV interaction after a pulse activation was inferred from a non-linear, single component regression. All numerical data processing and graph generation was performed using Excel 2011 (Microsoft) and Prism 7 (GraphPad software, inc.).

### Statistical analysis

All data were shown as means with SEM. Statistical significance as determined using two-tailed unpaired Student’s t-tests, Mann-Whitney U tests and the Wilcoxon matched-pairs signed rank test. Correlation coefficients between two data sets were calculated using Pearson *r* correlation tests or Spearman rank correlation tests. Data sets were assessed for their fit to a Gaussian distribution using the D’Agostino-Pearson omnibus K2 normality test prior to application of appropriate statistical test. A p-value of <0.05 was considered significant. *, P<0.05; **, P<0.01; ***, P<0.001; ****, P<0.0001. All statistical analyses were performed in Prism 7 (GraphPad software, inc.).

### Code availability

Our design algorithm is accessible via a web interface at http://104.131.81.59/, and the source code can be found at https://github.com/dannyhmg/germline.

### Data availability

The data that support the findings of this study are available from the corresponding author upon reasonable request.

## ACKNOWLEDGEMENTS

We thank S. Jonis, Y. Onderwater, H. Pires, P. van Bergeijk, and M. Harterink for reagents and L. Kapitein for technical advice. We also thank all the members of the Van den Heuvel, Akhmanova, Goldstein, Boxem, and Kapitein groups for helpful discussion and general support. We further thank A. Thomas for critically reading the manuscript. We acknowledge Wormbase and the Biology Imaging Center at the Faculty of Sciences, Department of Biology, Utrecht University. Some strains were provided by the Caenorhabditis Genetics Center (CGC), which is funded by NIH Office of Research Infrastructure Programs (P40OD010440). This work is part of program CW711.011.01 (S.v.d.H.) financed by the Netherlands Organization for Scientific Research (NWO) and was supported by a European Research Council (ERC) Synergy grant 609822 to A.A. D.J.D. was funded by a Helen Hay Whitney Foundation Postdoctoral fellowship and by NIH K99 GM115964. B.G. had grant support from NIH R01 GM083071.

## AUTHOR CONTRIBUTION

L.F., R.S., D.J.D., B.G., A.A., and S.v.d.H. designed the study and analyzed data. L.F. and S.v.d.H. wrote the manuscript. L.F. developed inducible knockout method and performed knockout experiments. L.F. and R.S. carried out optogenetic studies. D.J.D. developed germline-optimization algorithm and performed sequence optimization experiments.

## Supplementary Video Legends

Embryos are oriented with their anterior to the left in all movies.

All movies were made using spinning disk microscopy.

**Supplementary Video 1** Movie montage of mitosis in a one-cell *C. elegans* embryo expressing GFP::Tubulin (greyscale, microtubules), mCherry::TBG-1 (magenta, centrosomes) and mCherry::HIS-48 (magenta, DNA). Images, which are single planes, were made as a time-lapse with one acquisition per 2 seconds and played back at 10 frames per second, with time point 0 before the initiation of pronuclear meeting. Movie corresponds to the upper left panel in figure 1b.

**Supplementary Video 2** Movie montage of a mitotic spindle severing assay in a one-cell *C. elegans* embryo expressing GFP::Tubulin (greyscale, microtubules). The spindle is severed at the onset of anaphase using a pulsed UV laser (not visible), after which centrosomes are separated with speeds proportional to the net forces acting on them. Images, which are single planes, were made as a streaming acquisition with 0.5 seconds of exposure and played back at 10 frames per second, with time point 0 between late metaphase and anaphase initiation. Movie corresponds to figure 1c.

**Supplementary Video 3** Movie montage of a mitotic one-cell *C. elegans* embryo expressing diffuse cytosolic ePDZ::mCherry (greyscale, left; red, right) without PH::eGFP::LOV (green background, right). The movie shows diffuse localization of ePDZ::mCherry in presence of global and continual blue light exposure but in absence of a cortical LOV anchor. Images, which are single planes, were made as a time-lapse with one acquisition per 2 seconds for both 568 nm and 491 nm illumination and played back at 10 frames per second, with time point 0 starting at metaphase. The acquisition in the 568 nm channel at time point 0 shows localization of ePDZ::mCherry in complete absence of blue light, as embryos were kept in the dark before image acquisition. Movie corresponds to no main figure, and serves as a control for Supplementary Video 4.

**Supplementary Video 4** Movie montage of a mitotic one-cell *C. elegans* embryo expressing diffuse cytosolic ePDZ::mCherry (greyscale, left; red, right) and the membrane anchor PH::eGFP::LOV (green, right). The movie shows relocalization of diffuse ePDZ::mCherry to the cortex by global and continual activation of cortical LOV using blue light. Images, which are single planes, were made as a time-lapse with one acquisition per 2 seconds for both 568 nm and 491 nm illumination and played back at 10 frames per second, with time point 0 starting at late prophase. The acquisition in the 568 nm channel at time point 0 shows localization of ePDZ::mCherry in complete absence of blue light, as embryos were kept in the dark before image acquisition. Movie corresponds to the upper panels in figure 2b.

**Supplementary Video 5** Movie montage of a mitotic one-cell *C. elegans* embryo expressing diffuse cytosolic ePDZ::mCherry (inverted greyscale) and the membrane anchor PH::eGFP::LOV (not shown). The movie shows relocalization of diffuse ePDZ::mCherry to the posterior cortex by local activation of cortical LOV using low-intensity blue light. Activation of the ePDZ-LOV2 interaction is induced at the posterior cortex using local illumination with a 491 nm laser. The embryo was otherwise shielded from blue light before and during the experiment. Images, which are single planes, were made as a streaming acquisition with 0.5 seconds of exposure and played back at 10 frames per second, with time point 0 corresponding to late prophase. Movie corresponds to the lower panels in figure 2b.

**Supplementary Video 6** Movie montage of a four-cell *C. elegans* embryo expressing diffuse cytosolic ePDZ::mCherry (greyscale) and the membrane anchor PH::eGFP::LOV (not shown). The movie shows relocalization of diffuse ePDZ::mCherry to the cortex by activation of cortical LOV using a single pulse of blue light, and subsequent return to the dark state in absence of blue light. Images, which are single planes, were made as a time-lapse with one acquisition per 2 seconds played back at 10 frames per second, where time point 0 is the last acquisition before a single 1 second global pulse of 491 nm light. The acquisition at time point 0 shows localization of ePDZ::mCherry in complete absence of blue light, as the embryo was kept in the dark before and after global induction of the LOV-ePDZ interaction. Movie corresponds to figure 2c.

**Supplementary Video 7** Movie montage of a mitotic one-cell *C. elegans* embryo expressing endogenously labeled ePDZ::mCherry::RIC-8 (greyscale, left; red, right), the membrane anchor PH::eGFP::LOV and GFP::Tubulin (both green, right). The movie shows relocalization ePDZ::mCherry::RIC-8 to the cortex by global and continual activation of cortical LOV using blue light. Images, which are single planes, were made as a time-lapse with one acquisition per 2 seconds for both 568 nm and 491 nm illumination and played back at 10 frames per second, with time point 0 starting at late prophase. The acquisition in the 568 nm channel at time point 0 shows localization of ePDZ::mCherry::RIC-8 in complete absence of blue light, as embryos were kept in the dark before image acquisition. Movie corresponds figure 2g.

**Supplementary Video 8** Movie montage of a mitotic one-cell *C. elegans* embryo expressing endogenously labeled ePDZ::mCherry::GPR-1 (greyscale, left; red, right) in a ∆*gpr-2* genetic background, the membrane anchor PH::eGFP::LOV and GFP::Tubulin (both green, right). The movie shows relocalization of ePDZ::mCherry::GPR-1 to the cortex by global and continual activation of cortical LOV using blue light. Images, which are single planes, were made as a time-lapse with one acquisition per 2 seconds for both 568 nm and 491 nm illumination and played back at 10 frames per second, with time point 0 corresponding with early metaphase. The acquisition in the 568 nm channel at time point 0 shows localization of ePDZ::mCherry::GPR-1 in complete absence of blue light, as embryos were kept in the dark before image acquisition. Movie serves as a control to Supplementary Video 9.

**Supplementary Video 9** Movie montage of a mitotic one-cell *C. elegans* embryo treated with Gα RNAi expressing endogenously labeled ePDZ::mCherry::GPR-1 (greyscale, left; red, right) in a ∆*gpr-2* genetic background, the membrane anchor PH::eGFP::LOV and GFP::Tubulin (both green, right). The movie shows relocalization of ePDZ::mCherry::GPR-1 to the cortex by global and continual activation of cortical LOV using blue light. Images, which are single planes, were made as a time-lapse with one acquisition per 2 seconds for both 568 nm and 491 nm illumination and played back at 10 frames per second, with time point 0 corresponding with early metaphase. The acquisition in the 568 nm channel at time point 0 shows localization of ePDZ::mCherry::GPR-1 in complete absence of blue light, as embryos were kept in the dark before image acquisition. Movie corresponds to figure 3b.

**Supplementary Video 10** Movie montage of a mitotic one-cell *C. elegans* embryo expressing GFP::Tubulin (greyscale, microtubules) and PH::eGFP::LOV (greyscale, membrane). Invaginations (black arrows) are visible at the embryo membrane most pronouncedly in the posterior during late metaphase and anaphase. Images, which are single planes, were made as a streaming acquisition with 0.5 seconds of exposure and played back at 10 frames per second, with time point 0 between late metaphase and anaphase initiation. Movie corresponds to figure 3f.

**Supplementary Video 11** Movie montage of the subcortical area of a mitotic one-cell *C. elegans* embryo expressing GFP::Tubulin (inverted greyscale, microtubules), PH::eGFP::LOV (inverted greyscale, membrane). Invaginations are visible as dots protruding inwards from the embryo membrane during late metaphase and anaphase. Images, which are single planes, were made as a streaming acquisition with 0.25 seconds of exposure and played back at 10 frames per second, with time point 0 corresponding with anaphase, 50 seconds before telophase initiation. Movie corresponds to figure 3f.

**Supplementary Video 12** Movie montage of a mitotic one-cell *C. elegans* embryo expressing endogenously labeled ePDZ::mCherry::DHC-1 (greyscale, left; red, right), the membrane anchor PH::eGFP::LOV and GFP::Tubulin (both green, right). The movie shows relocalization of ePDZ::mCherry::DHC-1 to the cortex by global and continual activation of cortical LOV using blue light. Images, which are single planes, were made as a time-lapse with one acquisition per 2 seconds for both 568 nm and 491 nm illumination and played back at 10 frames per second, with time point 0 corresponding with early metaphase. The acquisition in the 568 nm channel at time point 0 shows localization of ePDZ::mCherry::GPR-1 in complete absence of blue light, as embryos were kept in the dark before image acquisition. Movie serves as a control to Supplementary Video 13.

**Supplementary Video 13** Movie montage of a mitotic one-cell *C. elegans* embryo treated with *gpr-1/2* RNAi expressing endogenously labeled ePDZ::mCherry::DHC-1 (greyscale, left; red, right), the membrane anchor PH::eGFP::LOV and GFP::Tubulin (both green, right). The movie shows relocalization of ePDZ::mCherry::DHC-1 to the cortex by global and continual activation of cortical LOV using blue light. Images, which are single planes, were made as a time-lapse with one acquisition per 2 seconds for both 568 nm and 491 nm illumination and played back at 10 frames per second, with time point 0 corresponding with early metaphase. The acquisition in the 568 nm channel at time point 0 shows localization of ePDZ::mCherry::DHC-1 in complete absence of blue light, as embryos were kept in the dark before image acquisition. Movie corresponds to figure 4b.

**Supplementary Video 14** Movie montage of a mitotic one-cell *C. elegans* embryo expressing endogenously labeled LIN-5::ePDZ::mCherry (greyscale, left; red, right) and the membrane anchor PH::eGFP::LOV and GFP::Tubulin (both green, right). The movie shows relocalization of LIN-5::ePDZ::mCherry to the cortex by global and continual activation of cortical LOV using blue light. Images, which are single planes, were made as a time-lapse with one acquisition per 2 seconds for both 568 nm and 491 nm illumination and played back at 10 frames per second, with time point 0 corresponding with metaphase. The acquisition in the 568 nm channel at time point 0 shows localization of LIN-5::ePDZ::mCherry in complete absence of blue light, as embryos were kept in the dark before image acquisition. Movie serves as a control to Supplementary Video 15.

**Supplementary Video 15** Movie montage of a mitotic one-cell *C. elegans* embryo treated with Gα RNAi expressing endogenously labeled LIN-5::ePDZ::mCherry (greyscale, left; red, right), the membrane anchor PH::eGFP::LOV and GFP::Tubulin (both green, right). The movie shows relocalization of LIN-5::ePDZ::mCherry to the cortex by global and continual activation of cortical LOV using blue light. Images, which are single planes, were made as a time-lapse with one acquisition per 2 seconds for both 568 nm and 491 nm illumination and played back at 10 frames per second, with time point 0 corresponding with metaphase. The acquisition in the 568 nm channel at time point 0 shows localization of LIN-5::ePDZ::mCherry in complete absence of blue light, as embryos were kept in the dark before image acquisition. Movie corresponds to figure 5b.

**Supplementary Video 16** Movie montage of a mitotic one-cell *C. elegans* embryo treated with *gpr-1/2* RNAi expressing endogenously labeled LIN-5::ePDZ::mCherry (not shown), the membrane anchor PH::eGFP::LOV and GFP::Tubulin (both inverted greyscale). The movie shows excessive rocking of centrosomes with associated pronuclei prior to mitotic spindle assembly. Images, which are single planes, were made as a streaming acquisition with 0.5 seconds of exposure and played back at 20 frames per second, with time point 0 corresponding with late prophase. Movie corresponds to no figure, but is discussed in the text.

**Supplementary Video 17** Movie montage of a mitotic one-cell *C. elegans* embryo expressing endogenously labeled LIN-5::ePDZ::mCherry (not shown) and the membrane anchor PH::eGFP::LOV (inverted greyscale). The movie shows separation of centrosomes and associated pronuclei in prophase upon global and continuous activation of LOV with blue light. Images, which are single planes, were made as a streaming acquisition with 0.5 seconds of exposure and played back at 10 frames per second, with time point 0 corresponding with prophase. Movie corresponds to no figure, but is discussed in the text.

**Supplementary Video 18** Movie montage of a mitotic one-cell *C. elegans* embryo treated with *gpr-1/2* RNAi expressing endogenously labeled LIN-5::ePDZ::mCherry (inverted greyscale) and the membrane anchor PH::eGFP::LOV (not shown). The movie shows anterior displacement of the spindle and subsequent inverted asymmetric division resulting in a small anterior and large posterior blastomere after local recruitment of LIN-5::ePDZ::mCherry to the anterior cortex. Images, which are single planes, were made as a time-lapse with one acquisition per 5 seconds and played back at 5 frames per second, with time point 0 corresponding with metaphase. Movie corresponds to figure 5g.

**Supplementary Video 19** Movie montage of a mitotic one-cell *C. elegans* embryo treated with *gpr-1/2* RNAi expressing endogenously labeled LIN-5::ePDZ::mCherry (inverted greyscale) and the membrane anchor PH::eGFP::LOV (not shown). The movie shows artificial transverse positioning of the metaphase mitotic spindle and its subsequent correction to an anterior-posterior position in late anaphase after local recruitment of LIN-5::ePDZ::mCherry to the opposing equatorial cortexes. Images, which are averages of groups of 2 subsequent frames, were made as a time-lapse with one acquisition per 2 seconds and played back at 5 frames per second, with time point 0 corresponding with metaphase. Movie corresponds to figure 5h.

**Supplementary Video 20** Movie montage of a mitotic two-cell *C. elegans* embryo treated with *gpr-1/2* RNAi expressing endogenously labeled LIN-5::ePDZ::mCherry (inverted greyscale) and the membrane anchor PH::eGFP::LOV (not shown). The movie shows artificial rotation of transverse aligned AB and P1 spindles to an anterior-posterior position, and concurrent reorientation of the cleavage planes after local recruitment of LIN-5::ePDZ::mCherry to the central region where AB and P1 cortexes touch. Images, which are averages of groups of 2 subsequent frames, were made as a streaming acquisition with 0.5 seconds of exposure and played back at 10 frames per second, with time point 0 corresponding with metaphase in the AB blastomere. Movie corresponds to figure 5i.

**Supplementary Video 21** Movie montage of a mitotic two-cell *C. elegans* embryo treated with *gpr-1/2* RNAi expressing endogenously labeled LIN-5::ePDZ::mCherry (inverted greyscale) and the membrane anchor PH::eGFP::LOV (not shown). The movie shows artificial rotation of transverse aligned AB and P1 spindles to an anterior-posterior position, and concurrent reorientation of the cleavage planes after local recruitment of LIN-5::ePDZ::mCherry to the central region where AB and P1 cortexes touch. Images, which are averages of groups of 2 subsequent frames, were made as a streaming acquisition with 0.5 seconds of exposure and played back at 20 frames per second, with time point 0 corresponding with metaphase in the AB blastomere. Movie corresponds to figure 5i.

